# Post-hoc analyses of Surrogate Markers of Non-Alcoholic Fatty Liver Disease (NAFLD) and Liver Fibrosis in Patients with Type 2 Diabetes in a Digitally-Supported Continuous Care Intervention: An Open Label, Non-Randomized, Controlled Study

**DOI:** 10.1101/293548

**Authors:** Eduardo Vilar-Gomez, Shaminie J. Athinarayanan, Rebecca N. AdamS, Sarah J. Hallberg, Nasir H. Bhanpuri, Amy L. McKenzie, Wayne W. Campbell, James P. McCarter, Stephen D. Phinney, Jeff S. Volek, Naga Chalasani

## Abstract

**Objective:** One-year of comprehensive continuous care intervention (CCI) through nutritional ketosis improves HbA1c, body weight and liver enzymes among type 2 diabetes (T2D) patients. Here, we report the effect of the CCI on surrogate scores of non-alcoholic fatty liver disease (NAFLD) and liver fibrosis.

**Methods:** This was a non-randomized longitudinal study, including adults with T2D who were self-enrolled to the CCI (n=262) or to receive usual care (UC, n=87) during one year. A NAFLD liver fat score [N-LFS] > −0.640 defined the presence of fatty liver. A NAFLD fibrosis score [NFS] of > 0.675 identified subjects with advanced fibrosis. Changes in N-LFS and NFS at one year were the main endpoints.

**Results:** At baseline, NAFLD was present in 95% of patients in the CCI and 90% of patients in the UC. At one year, weight loss of ≥ 5% was achieved in 79% of patients in the CCI vs. 19% of patients in UC (P<0.001). N-LFS mean score was reduced in the CCI group (−1.95±0.22, P<0.001) whereas it was not changed in the UC (0.47±0.41, P=0.26) (CCI vs. UC, P<0.001). NFS was reduced in the CCI group (−0.65±0.06, P<0.001) compared with UC (0.26±0.11, P=0.02) (P<0.001 between two groups). In the CCI group, the percentage of individuals with a low probability of advanced fibrosis increased from 18% at baseline to 33% at 1 year (P<0.001).

**Conclusions:** One year of a digitally-supported CCI significantly improved surrogates of NAFLD and advanced fibrosis in patients with type 2 diabetes.

**DATA SHARING:** Data sets and statistical code used for the current study are available from the corresponding author on reasonable request.

**Article Summary Strengths and limitations of this study:**

- This study highlights the beneficial effect of the CCI on NAFLD in high risk patients with T2D
- This study also identifies positive associations between glycemic improvements and improvements in ALT levels
- The assessment of resolution of steatosis and fibrosis is limited by the sensitivity and specificity of the non-invasive markers used in the study
- The patients were restricted in their carbohydrate intake and monitored for their nutritional ketosis state, but dietary energy, macronutrient and micronutrient intakes were not assessed.

## INTRODUCTION

Non-alcoholic fatty liver disease (NAFLD) is an important cause of chronic liver disease (CLD), hepatocellular carcinoma (HCC) and liver transplant worldwide, and is associated with increased risk of heart disease, diabetes, chronic kidney disease and malignancies[1–4]. NAFLD is highly prevalent (~70%) among patients with obesity and type 2 diabetes (T2D)[5]. T2D is usually associated with the more aggressive form of NAFLD, including non-alcoholic steatohepatitis (NASH, indicating significant hepatocellular injury) and advanced fibrosis[6] and is linked with high risk for all-cause and liver-related mortality[7–10]. Currently, there are no approved pharmacological interventions for NASH. Weight loss via lifestyle changes including dietary modification and exercise is the first-line intervention used in treating and improving NAFLD/NASH[11, 12]. However, the majority of patients do not achieve or sustain targeted weight loss goals[11, 13]. Previous studies show a close relationship between the degree of weight reduction and improvements in most of the NASH-related features, including steatosis, inflammation, fibrosis, insulin resistance and elevated liver enzymes, irrespective of the type of diet consumed[13–22]. However, there is an intense debate about what types of diet are most effective for treating NASH and, to date, the optimal degree of energy restriction and macronutrient composition of dietary interventions in subjects with NASH and T2D are not well defined[12].

Low-carbohydrate, high-fat (LCHF) and ketogenic diets have demonstrated a superior weight loss effect to low-fat, high-carbohydrate (LFHC) diets in adults with overweight and obesity[23–26] and short-term interventions with very low-carbohydrate diets are associated with improved insulin sensitivity and glycemic control[27, 28]. Lower consumption of carbohydrate, LCHF and ketogenic diets improve appetite control, satiety and/or reduce daily food intake helping to limit dietary energy consumption while maintaining patient-perceived vigor[29]. In patients with NAFLD, the beneficial effects of LCHF diets on liver enzymes and intrahepatic lipid content (IHLC) have been explored with contradictory results. Among studies with varied carbohydrate intakes, some reported a significant reduction of aminotransferases[16, 30–32], while others did not report significant changes in these enzymes[17, 33, 34]. A recent meta-analysis of pooled data from 10 clinical trials reported that low carbohydrate diet (LCD) in patients with NAFLD led to a significant reduction in IHLC[35].

We recently demonstrated that one-year of a telemedicine-based comprehensive continuous care intervention (CCI) with carbohydrate restriction-induced ketosis and behavior change support significantly reduced HbA1c level and medication usage in patients with T2D[36]. The effectiveness of the CCI relies in maintaining a carbohydrate-restricted diet, and monitoring compliance with the dietary regimen by assessing the patient’s nutritional ketosis by blood tests during the year. We also demonstrated that one year of the CCI was effective in improving liver enzymes, where mean alanine aminotransferase (ALT), aspartate aminotransferase (AST) and alkaline phosphatase (ALP) were reduced by 29%, 20% and 13%, all P<.01, respectively. These findings not only highlight the beneficial effect of the CCI on diabetes management, but also in ameliorating the liver-related injury. These changes were not reported in the usual care (UC) patients receiving standard diabetes care treatment. Therefore, in the current post-hoc analysis, we assessed one-year within- and between-group (CCI vs. usual care; UC) differences in non-invasive liver markers of steatosis (NAFLD liver fat score) and fibrosis (NAFLD fibrosis score) in the full study sample (CCI and UC cohorts). In addition, we assessed these outcomes, in the subgroup of patients with abnormal ALT at baseline (ALT levels of > 30 U/L in men and >19 U/L in women). Among all patients, ancillary aims included assessing if changes in weight and HbA1c were associated with ALT and metabolic parameter improvements, and potential relationships between changes in the ALT with other metabolic parameters.

## METHODS

The design and primary results of this study were previously published, and the current results are based on a 1-year post-hoc analysis using the data collected from the same cohort in that clinical study (*Clinical trials.gov identifier: NCT02519309*)[36]. A brief description of the study design, participants and interventions are listed in the **supplementary appendix (Methods section)**. Briefly, this was a non-randomized and open-label controlled longitudinal study, including patients 21 to 65 years of age with a diagnosis of T2D and a BMI > 25 kg/m^2^. Further, patients were excluded if they had significant alcohol intake (average consumption of three or more alcohol-containing beverages daily or consumption of more than 14 standard drinks per week), presence of any other cause of liver disease or secondary causes of NAFLD and decompensated cirrhosis.

### Patient and public involvement

Patients were not involved in the design and implementation of the study. Patient participants have been thanked for their participation in all resulting manuscripts and will receive information on publications upon study completion.

### Study Recruitment and intervention

Patients participating in the CCI had access to a remote care team consisting of a personal health coach and medical providers (physician or nurse practitioner). The participants in the CCI self-selected between two different educational modes; either via on-site education classes (n=136, CCI-onsite) or via web-based educational content (n=126, CCI-virtual). The CCI patients were routinely assessed for nutritional ketosis based on blood beta-hydroxybutyrate (BHB) concentrations. We also recruited and followed a cohort of UC patients with T2D (n=87) who received a standard diabetes care treatment from their primary care physician or endocrinologist without modification[36,37].

### Outcomes Primary outcomes-NAFLD liver fat and liver fibrosis by non-invasive surrogate markers

NAFLD liver fat score (N-LFS) is a surrogate marker of fatty liver which includes the presence of the metabolic syndrome, type 2 diabetes, fasting serum insulin, AST and the AST/ALT ratio. An N-LFS cutoff of > −0.640 predicts liver fat (> 5.56 % of hepatocytes) with a sensitivity of 86% and specificity of 71% [38, 39]. NAFLD fibrosis score (NFS) is a widely validated biomarker for identifying patients at different risks of fibrosis severity. NFS is derived from age, BMI, hyperglycemia, the AST/ALT ratio, platelet and albumin. The NFS threshold of < −1.455 can reliably exclude patients with advanced fibrosis (negative predictive value ≈ 92%) and > 0.675 can accurately detect subjects with advanced fibrosis (positive predictive value ≈ 85%)[40–42]. The equations for calculating both scores are displayed in the **supplementary appendix (Methods section)**.

### Ancillary outcomes-other biochemical markers

Results from other metabolic (HbA1c, fasting glucose, fasting insulin, HOMA-IR, triglycerides, total cholesterol, HDL cholesterol and LDL cholesterol), liver (ALT, AST, ALP), kidney (creatinine, eGFR), beta-hydroxybutryrate (BHB) and high sensitivity C-reactive protein (hsCRP) parameters were previously published in the full CCI and UC cohort [36]. These additional biochemical markers were assessed in the subset analyses of patients with abnormal ALT at baseline[43].

### Statistical analyses

First, we examined the assumptions of normality and linearity. According to Kline’s (2011) guidelines [44], seven outcomes (i.e., N-LFS, ALT, AST, fasting insulin, triglycerides, c-reactive protein, beta hydroxybutyrate) were positively skewed. We explored two approaches to handling the skewed variables: natural log-transformations and removing the top 1% of values. For N-LFS which includes both positive and negative values, a modulus log-transformation[45] was performed instead of a natural log-transformation. For every variable except triglycerides, both approaches resulted in new skew and kurtosis values falling within the acceptable range.

We conducted sensitivity analyses related to our first aim to compare the two approaches. The results did not differ between the two approaches, and to make interpretation feasible, we report results from the approach of removing the top 1% of values for the LMM analyses. For triglycerides, analyses were performed on the log-transformed variable; p-values reported are based on analyses with the transformed variable but the means and standard errors reported were computed from the original variable without any adjustments. For both ANCOVA and correlation analyses, the natural or modulus log-transformed variables were used to determine the association.

The first aim of the study was to examine (1) within-group changes in the study outcomes from baseline to 1 year, and (2) between-group differences (CCI vs. UC) in the study outcomes at 1 year. The on-site and virtual CCI patients were grouped together for analyses since no significant differences were observed in biochemical markers between these two modes of educational delivery[36]. We performed linear mixed-effects models (LMMs) in SPSS statistics software to estimate the within- and between-group differences. The LMMs included fixed effects for time, group (CCI vs. UC), and time by group interaction. Covariates included baseline age, sex, race (African American vs. other), diabetes duration, body mass index (BMI), and insulin use. This maximum likelihood-based approach uses all available repeated data, resulting in an intent-to-treat analysis. An unstructured covariance structure was specified for all models to account for correlations between repeated measures. Most analyses were conducted on a subsample of participants with abnormal (>30 U/L in men and >19 U/L in women) [46] ALT at baseline (195 of 347; 157 CCI and 38 UC). We also conducted analyses assessing changes in N-LFS, NFS, albumin, and platelets on the full study sample because results were not previously reported. In addition, we examined changes in the proportions of participants meeting clinically-relevant cut-offs for N-LFS, NFS, and ALT. Within-group changes in the proportions from baseline to 1 year were assessed using McNemar’s test. Between-group differences in proportions were assessed using Chi-Square test. For this set of analyses, multiple imputation (20 imputations) was used to replace missing values from baseline and 1 year with a set of plausible values, facilitating an intent-to-treat analysis.

The second study aim was to explore relationships between (1) changes in weight loss and HbA1c categories and its associations with ALT and metabolic parameters improvements and (2) changes in ALT and metabolic variables. Multiple imputation was also used to handle missing data for aim 2 analyses. We performed one-way longitudinal ANCOVA analyses for comparisons between different cutoffs of weight loss (<5%, 5-10% and >10%) and with changes in diabetes- and liver-related continuous variables. Covariates included baseline value of the dependent variables and body mass index (BMI). Trend analyses were performed using Mantel-Haenszel χ^2^ tests to assess changes in the proportions of patients meeting clinical cut-offs (for ALT, N-LFS and NFS normalization) within different weight and HbA1c categories. An adjusted odds ratio was calculated to measure the strength of association between HbA1c changes and ALT normalization using logistic regression (Figure 1B). The logistic regression analysis was adjusted by BMI, age, gender and baseline dependent covariates. Unadjusted and adjusted Pearsons’ correlations were performed to identify relationships between changes in ALT levels and changes in metabolic- and lipid-related parameters from baseline to 1 year. Adjusted correlations were also performed while controlling for baseline dependent covariates, baseline age, sex, race (African American vs. other), diabetes duration, body mass index (BMI), and insulin use. All confidence intervals, significance tests, and resulting *P* values were two-sided, with an alpha level of 0.05. A Bonferroni correction was applied to each set of analyses (LMM or ANCOVA) to control the family-wise error rate (FWER). The Bonferroni adjusted p-value =0.05/19 variables = 0.0025 was used to determine statistical significance for each set of hypothesis driven analyses.

**Figure 1.**
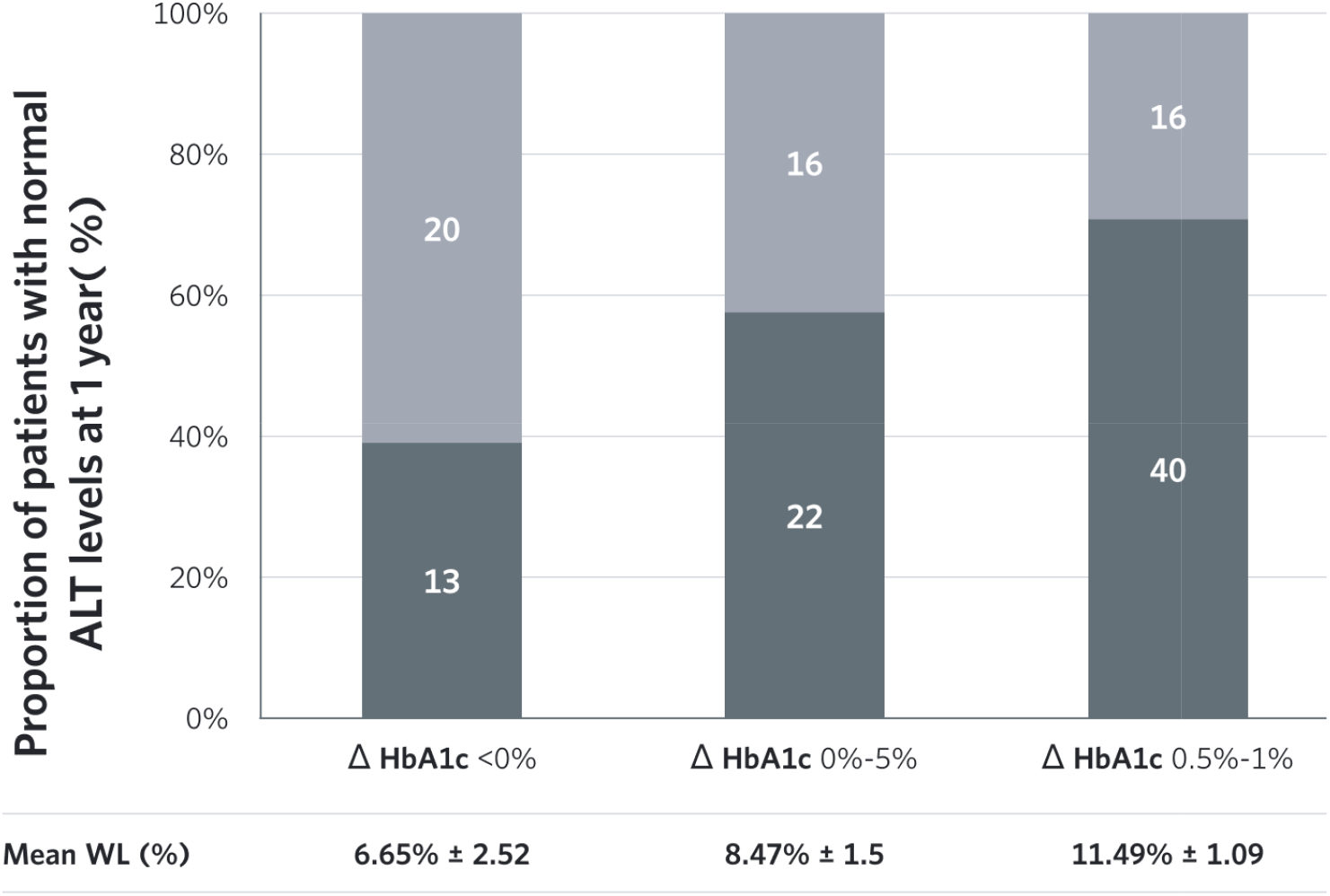
Association between reduction in HbA1c (%) and normalization of ALT* levels at 1 year of intervention in CCI gro ull CCI cohort (n=272) proportion of patients with ALT normalization were observed in HbA1c (%) reduction categories 0.5-10%; 71% and >

**Figure 2.**
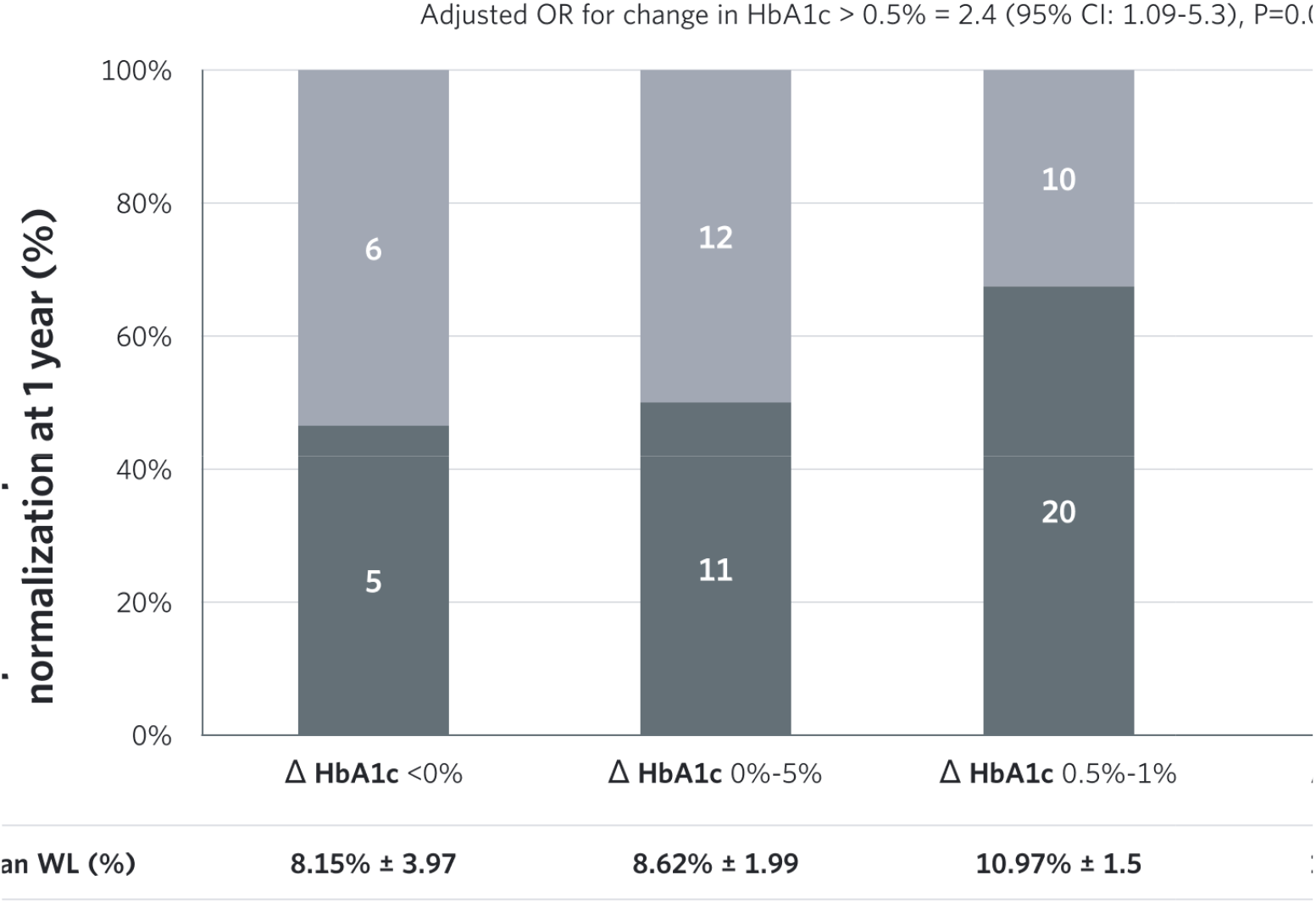
CCI patients with increased levels of ALT at baseline (n=153) ïr proportion of patients with ALT normalization were observed in HbA1c (%) reduction categories 0.5-1%; 67% and ≥1; ted OR for change in HbA1c > 0.5% = 2.4 (95% CI: 1.09-5.3), P=0.029

**Figure 3.**
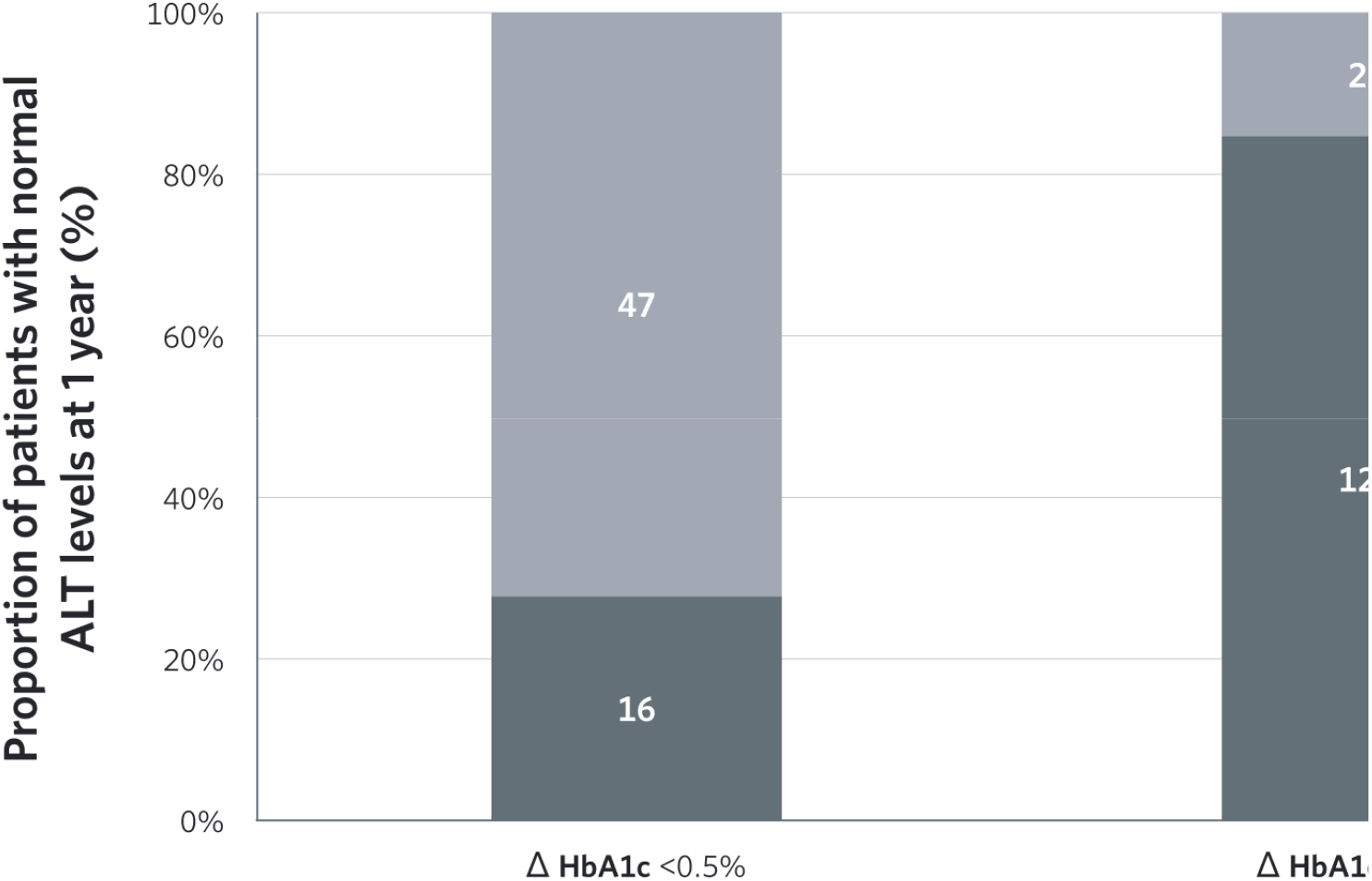
CCI patients with weight loss ≥5% (n=207). patients with weight loss ≥ 5%, higher levels of ALT normalization (85%) were observed in patients with HbA1c (%) .5%.

**Figure 4.**
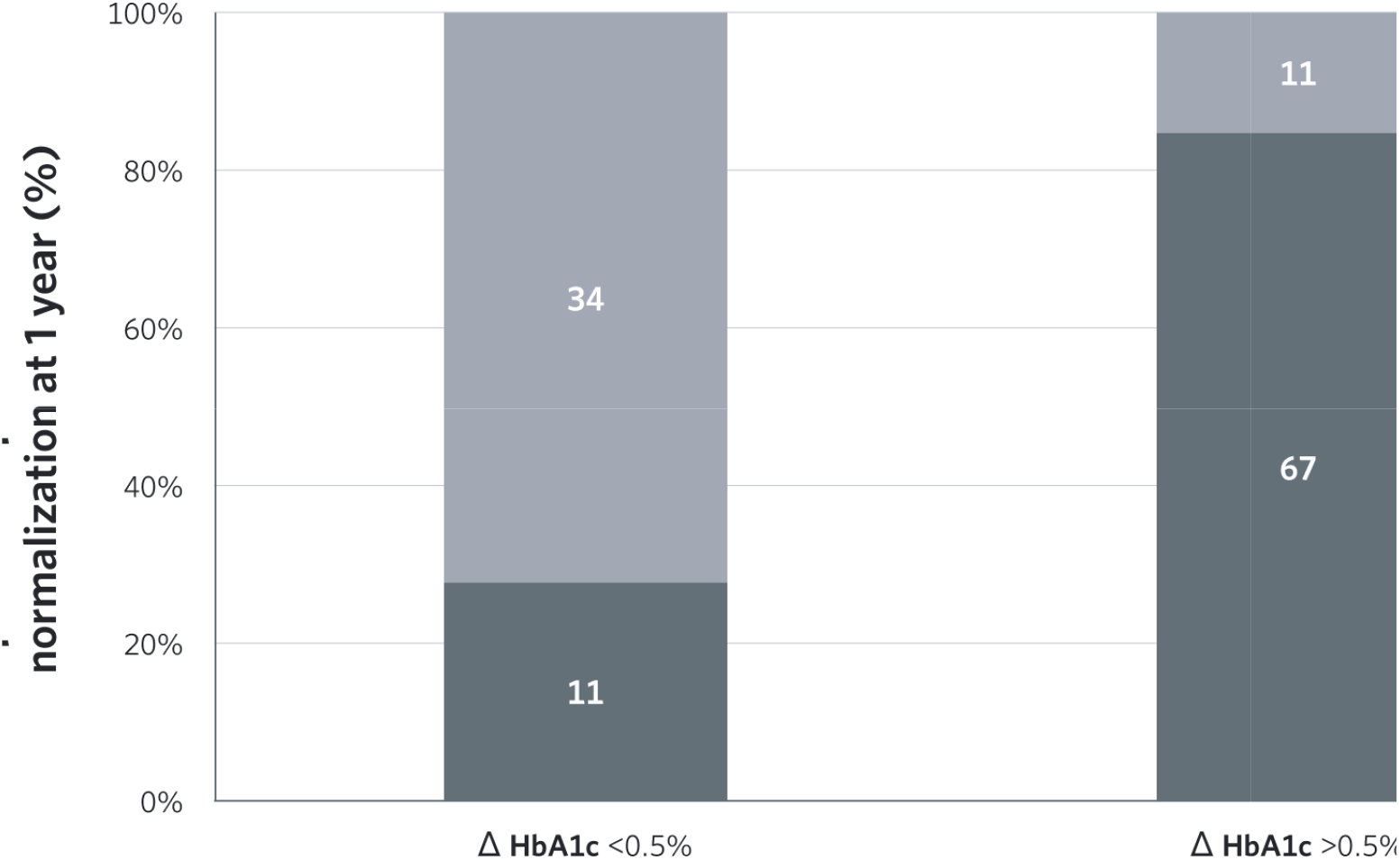
CCI patients with increased levels of ALT at baseline and weight loss ≥ 5% (n=123). patients with weight loss ≥ 5% and abnormal ALT levels at baseline, higher levels of ALT normalization (86%) were o ients with HbA1c (%) reduction of ≥0.5%. ALT levels ≤ 19 in women and ≤ 30 in men.

## RESULTS

### Baseline features of participants

Recruitment and baseline results were published previously[36]. Briefly, between August 2015 and April 2016, 262 and 87 patients were enrolled in the CCI and UC groups, respectively. **Supplemental Figure 1** shows the flow of patients through the study. At baseline, average age was 53.4 ± 8.7 years and 226 participants (65%) were female. The average time since T2D diagnosis was 8.3 ± 7.2 years and 314 subjects (90%) were obese with a mean BMI of 39.5[35]. Two-hundred and ninety-three participants (84%) were on medication for diabetes and 118 (34%) were insulin users[36]. The proportion of patients with abnormal ALT was higher in CCI (58%) compared to the UC (44%). At baseline, 330 subjects (95%) had suspicion of NAFLD and fewer patients (69 of 349 [20%]) had a NFS threshold of < −1.455 indicating low probability of advanced fibrosis. Compared to UC, mean baseline BMI was significantly higher in patients in the CCI. The remaining patient demographics and baseline features were generally not different between the two groups [36, 47].

### Influence of intervention and time on 1-year study endpoints Non-invasive markers of steatosis (N-LFS) and NAFLD fibrosis (NFS)

After one year, the CCI decreased N-LFS and NFS for the full cohort and among patients with abnormal ALT at baseline, whereas no changes were observed in the UC full cohort or subset (Table 1). There were significant between group (CCI vs. UC) differences in N-LFS and NFS observed in both the full and abnormal baseline ALT cohort at one year (Table 1). Notably, the proportion of patients with suspected steatosis reduced from 95% to 75% at 1 year in the CCI whereas no change occurred in UC. At 1 year, the proportion of patients without fibrosis increased from 18% to 33% in CCI group, P<0.001, but no change occurred in the UC. Similar to the full cohort, the proportion of patients with suspected steatosis was reduced from 99% to 76%, P<0.001 and proportion of those without fibrosis increased from 20% to 37%, P<0.001 through one year among CCI patients with abnormal ALT levels (Table 2). Between-group (CCI vs. UC) differences at 1 year are listed in Table 1.

### Metabolic parameters

At 1 year, beneficial changes observed in the metabolic parameters of the full CCI cohort[35,44] were also reported in the subset of patients with abnormal baseline ALT, including reduction of HbA1c, fasting glucose, fasting insulin, HOMA-IR, triglycerides (All, P<0.001), and increase of HDL-C (P<0.001) (Table 1). No changes in metabolic parameters were observed in the UC group. Between-group (CCI vs. UC) differences at 1 year are listed in Table 1.

### Other liver-related, kidney-function tests and parameters

Among CCI patients with abnormal ALT at baseline, significant reductions in the liver enzymes were observed (Table 1), as previously reported in the full CCI cohort. No changes in liver-related tests were observed in the UC group. Among patients with increased ALT levels at baseline, 93 (61%) of 153 participants enrolled in the CCI vs. 3 (8%) of 38 patients in UC had ALT normalization at 1 year (Table 2). Significant within-CCI changes were observed for albumin and platelet in the full CCI cohort, whereas in the subsample of patients with abnormal baseline ALT, there was only a significant decrease in the platelet (Table 1). As reported in the full CCI cohort [35], significant changes in c-reactive and beta-hydroxybutyrate concentrations were found in the subset of CCI patients with abnormal baseline ALT over 1 year. These changes were not found in the UC group. When adjusted for multiple comparisons, no significant changes in creatinine or eGFR were found in either the CCI or UC group. Between-group differences at 1 year are listed in Table 1.

**Table 1.**
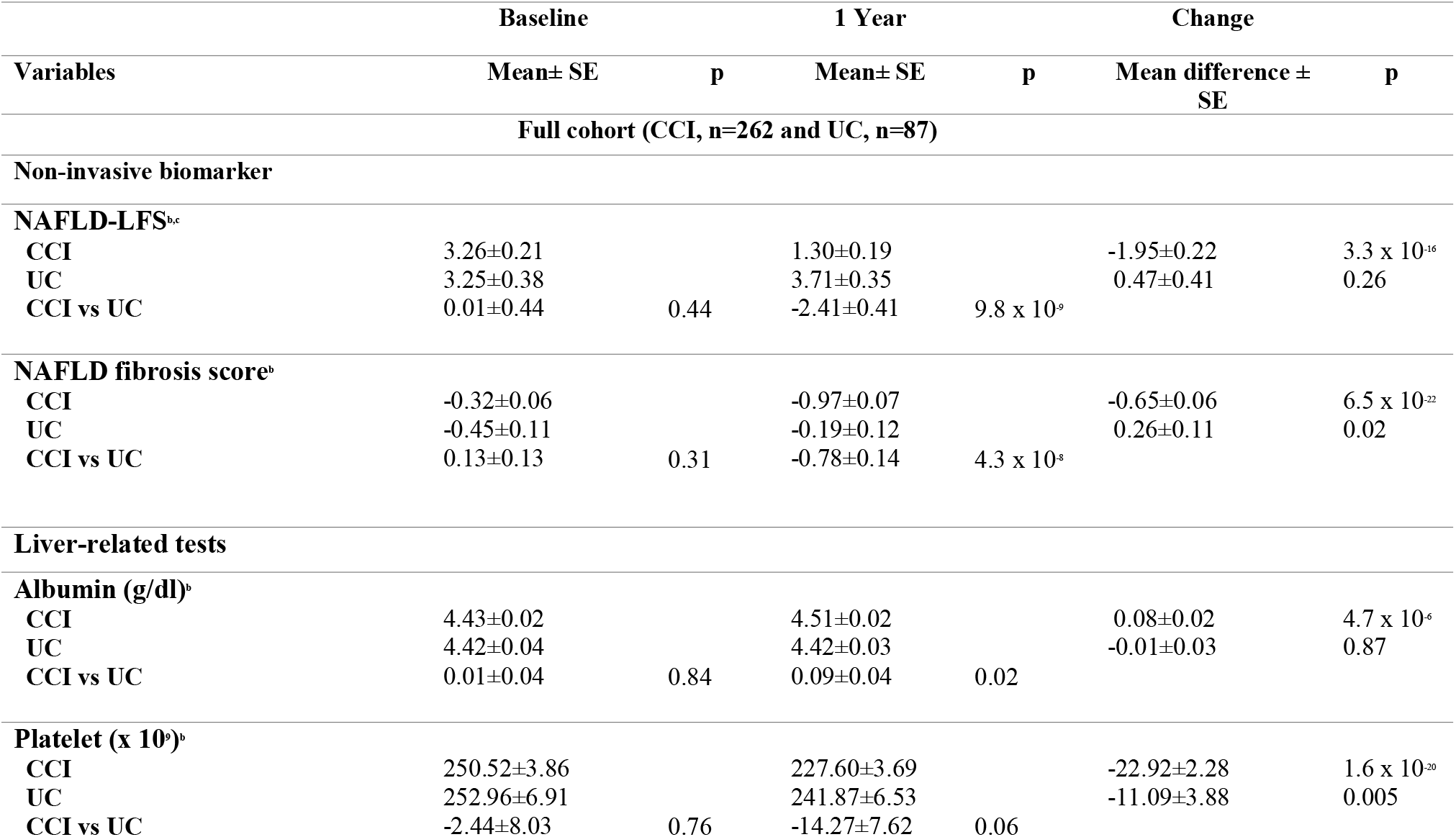
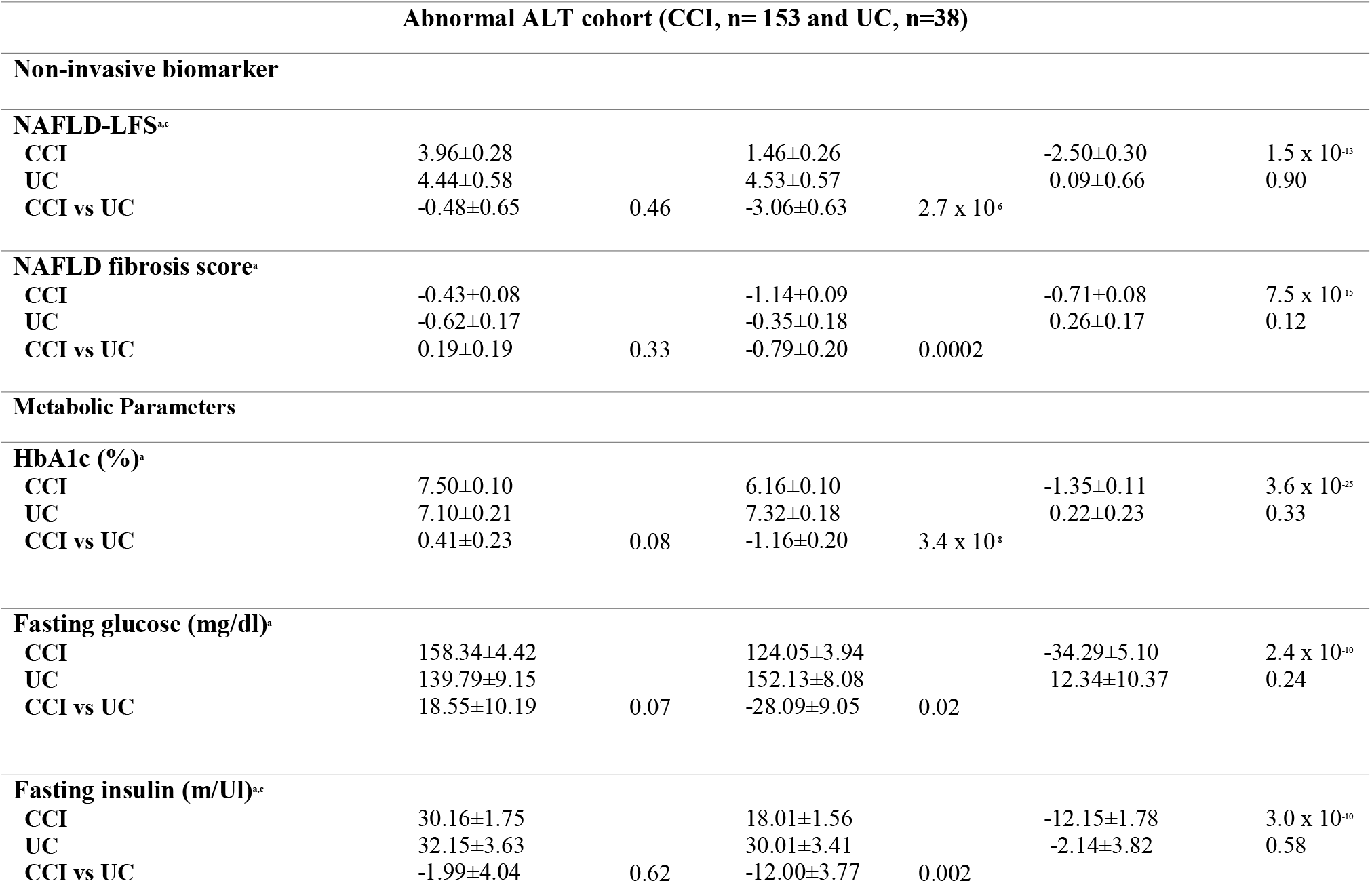
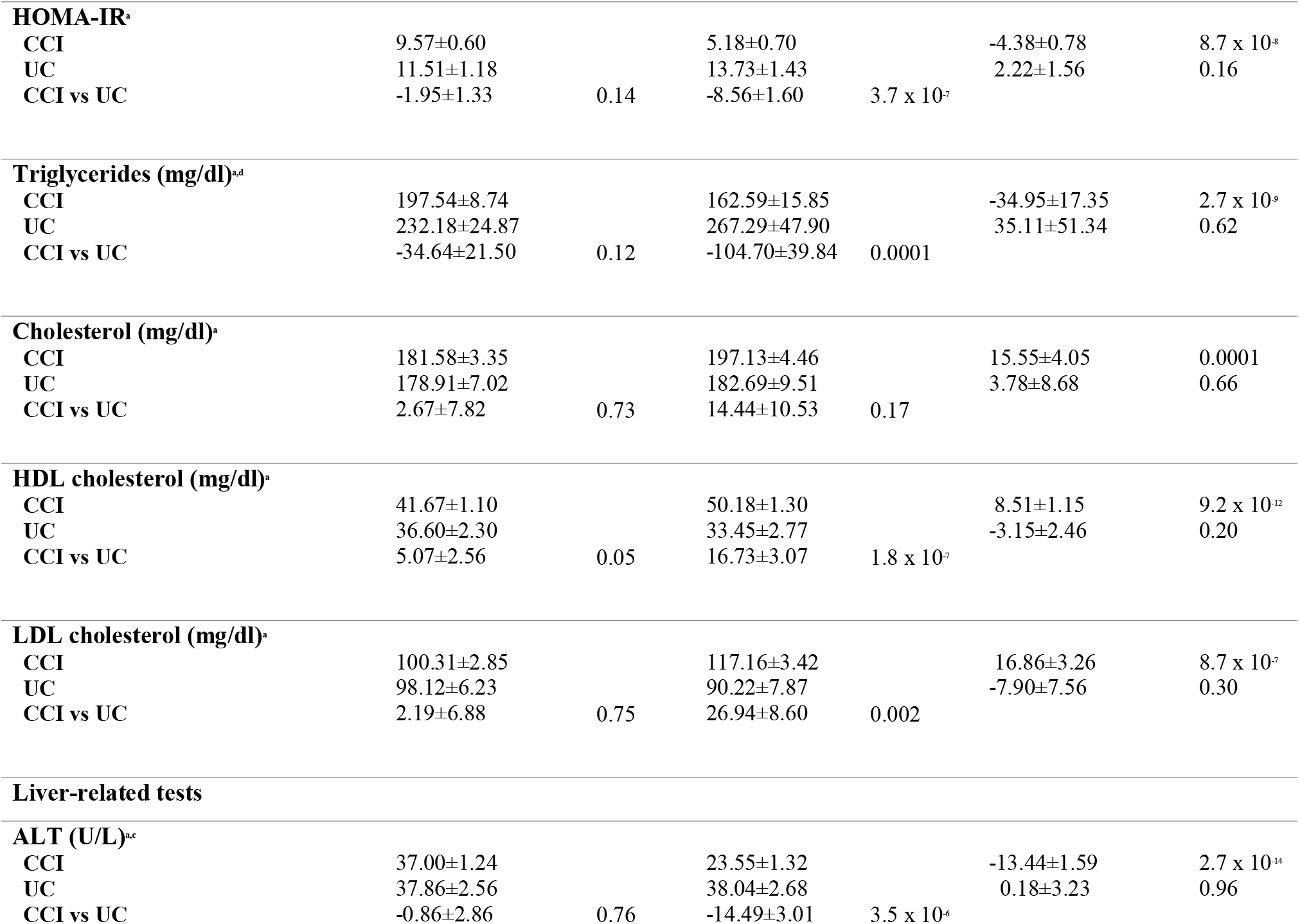
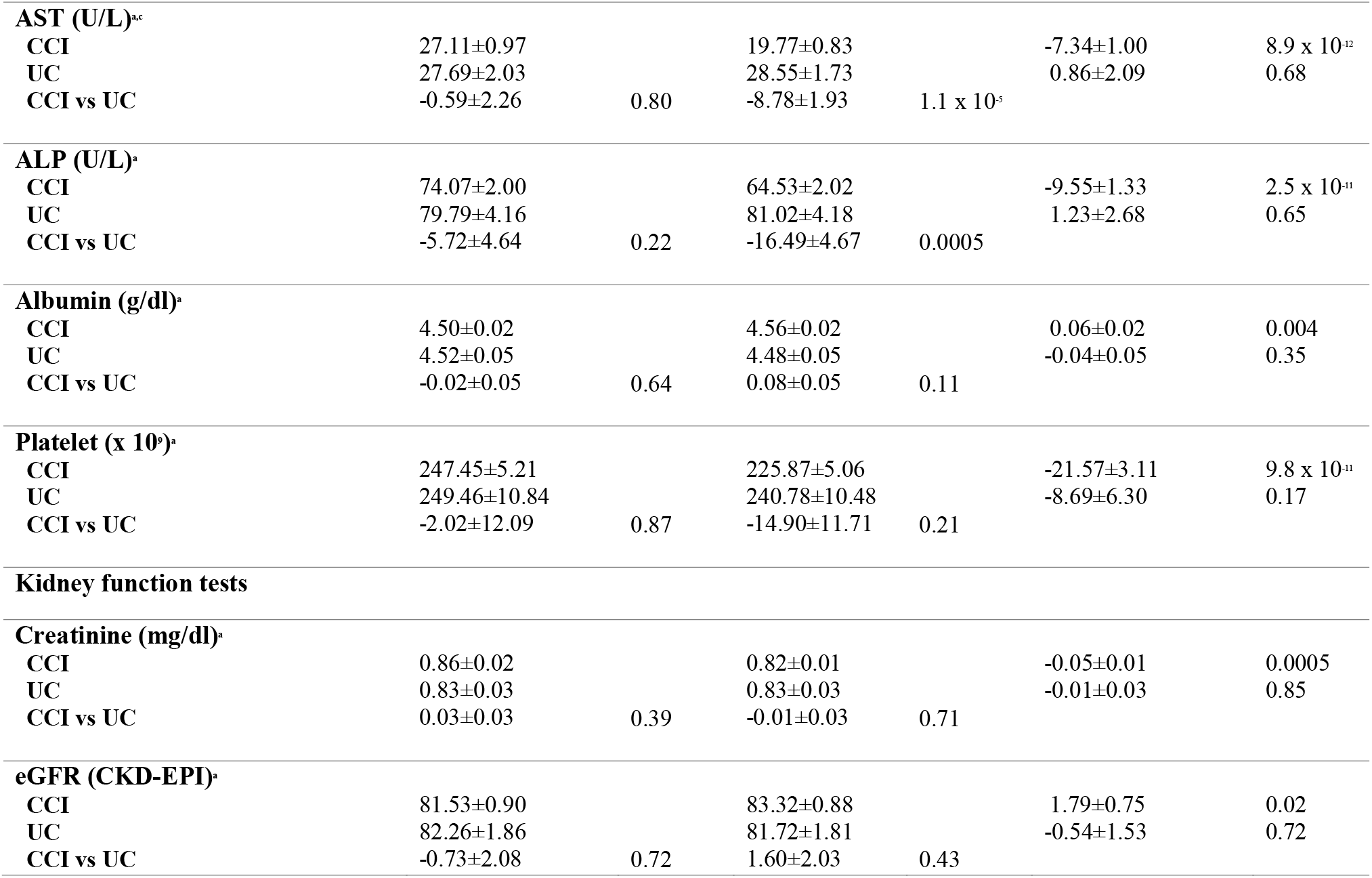
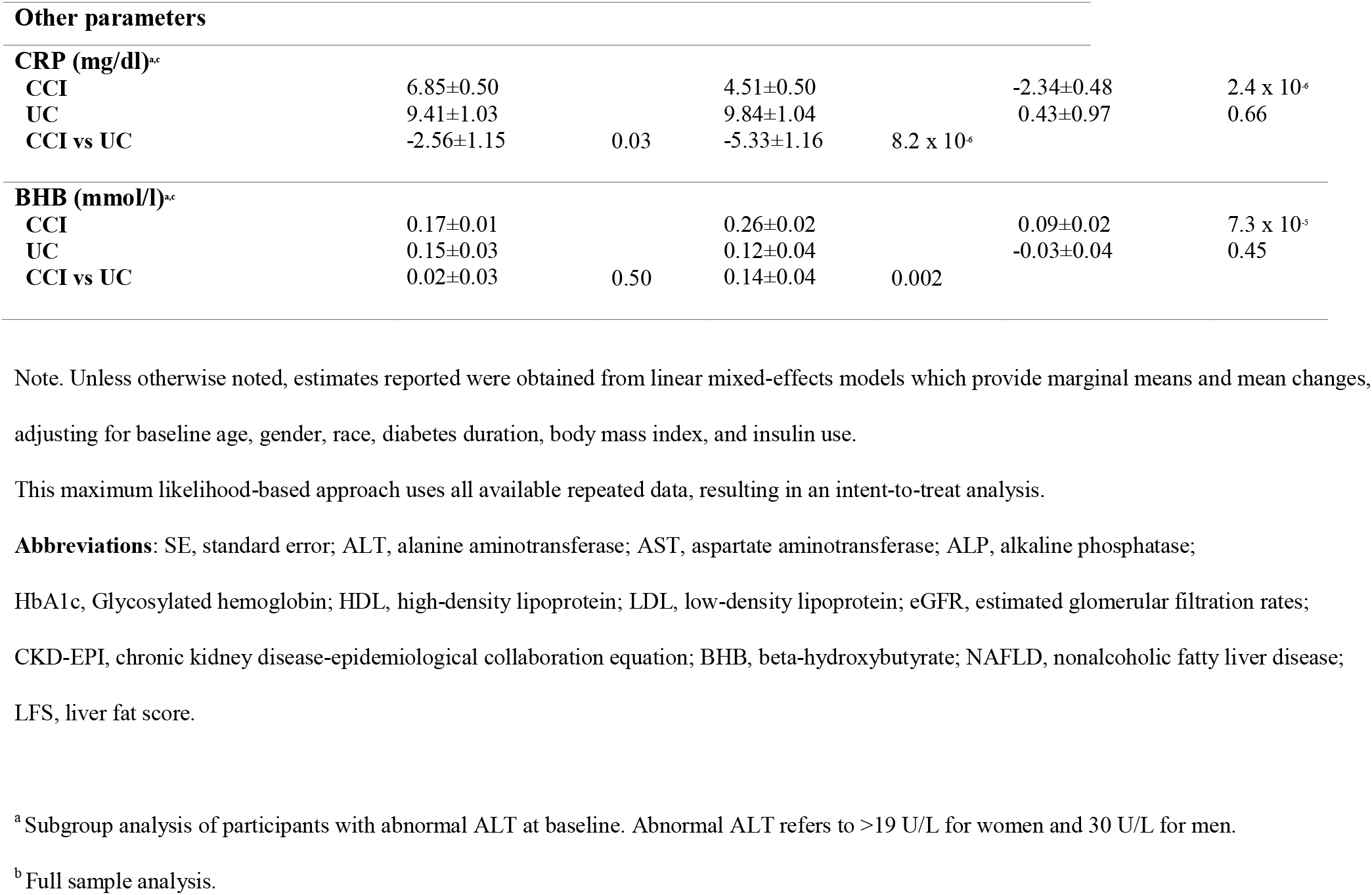
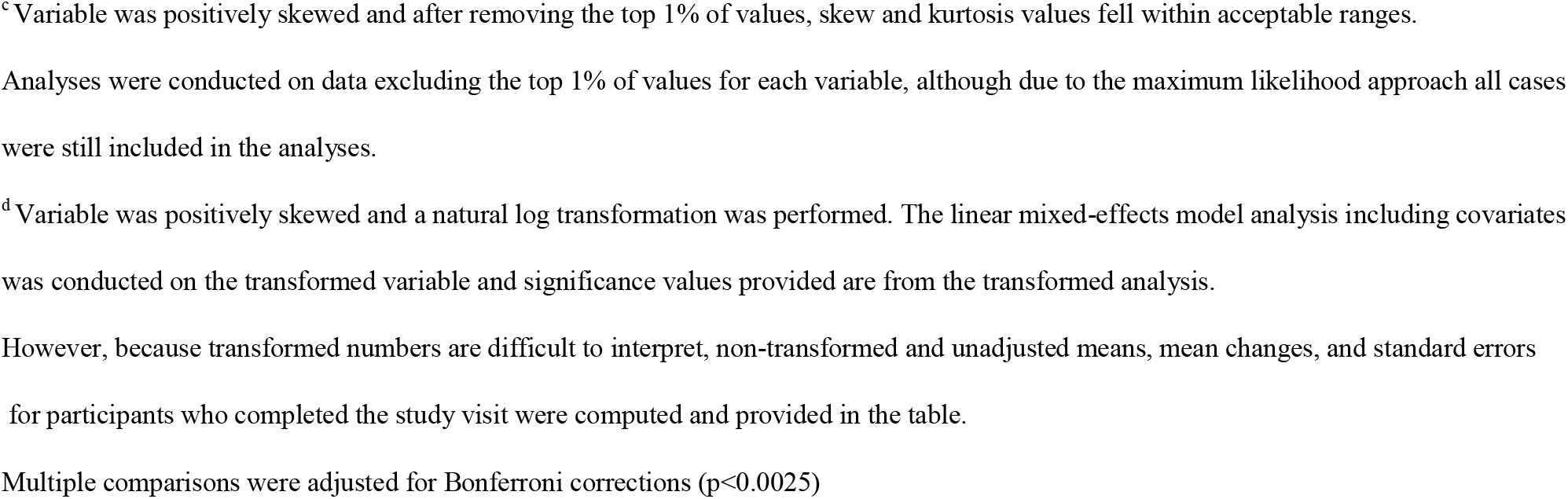
Estimated marginal means and mean changes in metabolic, liver-related and non-invasive markers at baseline and after one year of the CCI and UC interventions.

**Table 2.** Resolution of abnormal ALT, steatosis and fibrosis (as estimated using non-invasive liver markers cut-off) from baseline to one year in CCI and UC

| Variables | Continuous Care Intervention |  |  | Usual Care |  |  | Between-groups<br>P values<sup>b</sup> |
| --- | --- | --- | --- | --- | --- | --- | --- |
|  | Baseline | 1 year | P value<sup>a</sup> | Baseline | 1 year | P value<sup>a</sup> |  |
| **Full Cohort** |  | **n=262** |  |  | **n=87** |  |  |
| Abnormal ALT, n (%) † | 153 (58%) | 60 (23%) | 8.1x10<sup>-11</sup> | 38 (44%) | 35 (40%) | .664 | 0.006 |
| NAFLD-LFS |  |  |  |  |  |  |  |
| > -0.640 | 250 (95%) | 197 (75%) | 7.9x10<sup>-10</sup> | 80 (92%) | 79 (91%) | .678 | 0.002 |
| NAFLD fibrosis score |  |  |  |  |  |  |  |
| < -1.455 | 46 (18%) | 87 (33%) | 3.9x10<sup>-7</sup> | 23 (26%) | 22 (25%) | 1.0 | 0.139 |
| **Abnormal ALT at baseline** |  | **n=153** |  |  | **n=38** |  |  |
| NAFLD-LFS |  |  |  |  |  |  |  |
| > -0.640 | 151 (99%) | 117 (76%) | 1.8x10<sup>-7</sup> | 35 (92%) | 37 (97%) | 0.625 | 0.007 |
| NAFLD fibrosis score |  |  |  |  |  |  |  |
| < -1.455 | 30 (20%) | 56 (37%) | 4.1x10<sup>-5</sup> | 11 (29%) | 11 (29%) | 1.0 | 0.266 |

NAFLD-LFS cutoff > -0.640 for detecting liver fat > 5.56 % (sensitivity 86% and specificity 71%).
NAFLD fibrosis score < -1.455 corresponds with low probability of advanced fibrosis (NPV ≈ 92%) and > 0.675 indicates high probability of advanced fibrosis (PPV ≈ 85%).
† Abnormal ALT refers to >19 U/L for women and 30 U/L for men.
<sup>a</sup>McNemar's or <sup>b</sup>Chi-square tests were used when appropriated

### Associations between weight loss and study outcomes in the CCI group

At one year, weight loss of ≥ 5% was achieved in 79% of CCI patients with 54% achieving weight loss of ≥ 10%. The proportion of patients losing weight was lower in the UC group with only 17 UC participants (19.5%) achieving ≥5% weight loss and only 4 (6%) with ≥10% weight loss (Supplementary Figure 2). In the CCI group, there was a trend toward greater mean percentage weight loss (WL) by higher baseline BMI classification, especially in patients losing more than 5% or 10% of body weight (Supplementary Table 1). As shown in Table 3, there were relationship trends between the degree of 1-year of WL (%) and changes in liver, metabolic and non-invasive markers of steatosis and fibrosis among CCI participants. At 1 year, the CCI patients who achieved WL ≥ 10% showed the greatest reductions in N-LFS (P<0.001) and NFS (P<0.001), whereas no statistically significant differences were found between patients with WL from 5%-10% versus <5%. Similarly, patients who achieved WL ≥ 10% also showed decreases in HbA1c (P<0.001) and triglycerides (P<0.001) from baseline to 1 year. The one-year probability of suspected fatty liver (N-LFS ≥-0.64) was lower (66%) among patients with WL ≥ 10% compared to the other WL groups (<5% [85%] and 5%-10% [86%]). The proportion of patients with low likelihood of fibrosis at 1 year was higher among patients with WL ≥ 10% (41%) vs. patients with WL of 5-10% (26%) and <5% (22%).

### Correlation analyses between changes in ALT levels with changes in metabolic parameters in the CCI group

In the CCI group, changes in HbA1c, weight and fasting glucose from baseline to 1 year were associated with changes in ALT levels in the full cohort (HbA1c, r=0.148, P=0.03; weight, r=0.198, P=0.004; fasting glucose, r=0.176, P=0.004), and among patients with abnormal levels of ALT at baseline (HbA1c, r=0.253, P=0.005; weight, r=0.278, P=0.003, fasting glucose, r=0.305, P<0.001) (Table 4). Changes in other lipid markers did not correlate with changes in ALT levels (Table 4). Figures 1A-1D displays 1-year associations between change in HbA1c and normalization of ALT levels. In the full CCI group, 141 (70%) of 201 patients with HbA1c reductions of ≥ 0.5% at 1 year had normal ALT levels (Figure 1A). Among CCI patients with abnormal ALT levels at baseline, 77 (65%) of 119 patients with a reduction of ≥ 0.5% in HbA1c showed normalization of ALT levels (Figure 1B). One-year reduction of ≥ 0.5% in HbA1c increased the odds of ALT normalization 2.4 fold (95% CI: 1.09-5.3) after controlling for baseline levels of HbA1c, BMI, ALT, diabetes duration, insulin use and weight loss (%) at 1 year. Given that weight reductions (≥ 5%) can be associated with changes in HbA1c level, we sought to explore whether a reduction of ≥ 0.5% in HbA1c was still associated with ALT normalization, independent of weight loss (≥ 5%) (Figures 1C-D). A reduction of ≥ 0.5% in HbA1c was associated with higher rates of ALT normalization, regardless of whether or not 5% weight loss was achieved, P<.001.

### Safety

Adverse events during this trial were previously reported[35]. Mean platelet count was reduced in the CCI (−22.9 ± 2.3, P<0.001) vs. UC group (−11.1 ± 3.9, P=0.005); however, the proportion of patients with a platelet count below 150 × 10^9^ L was not different between groups. There was no hepatic decompensation (variceal hemorrhage, ascites or hepatic encephalopathy) or ALT flare-up (≥5 times the upper limit of normal) reported during the trial in either the CCI or UC group.

**Table 3.**
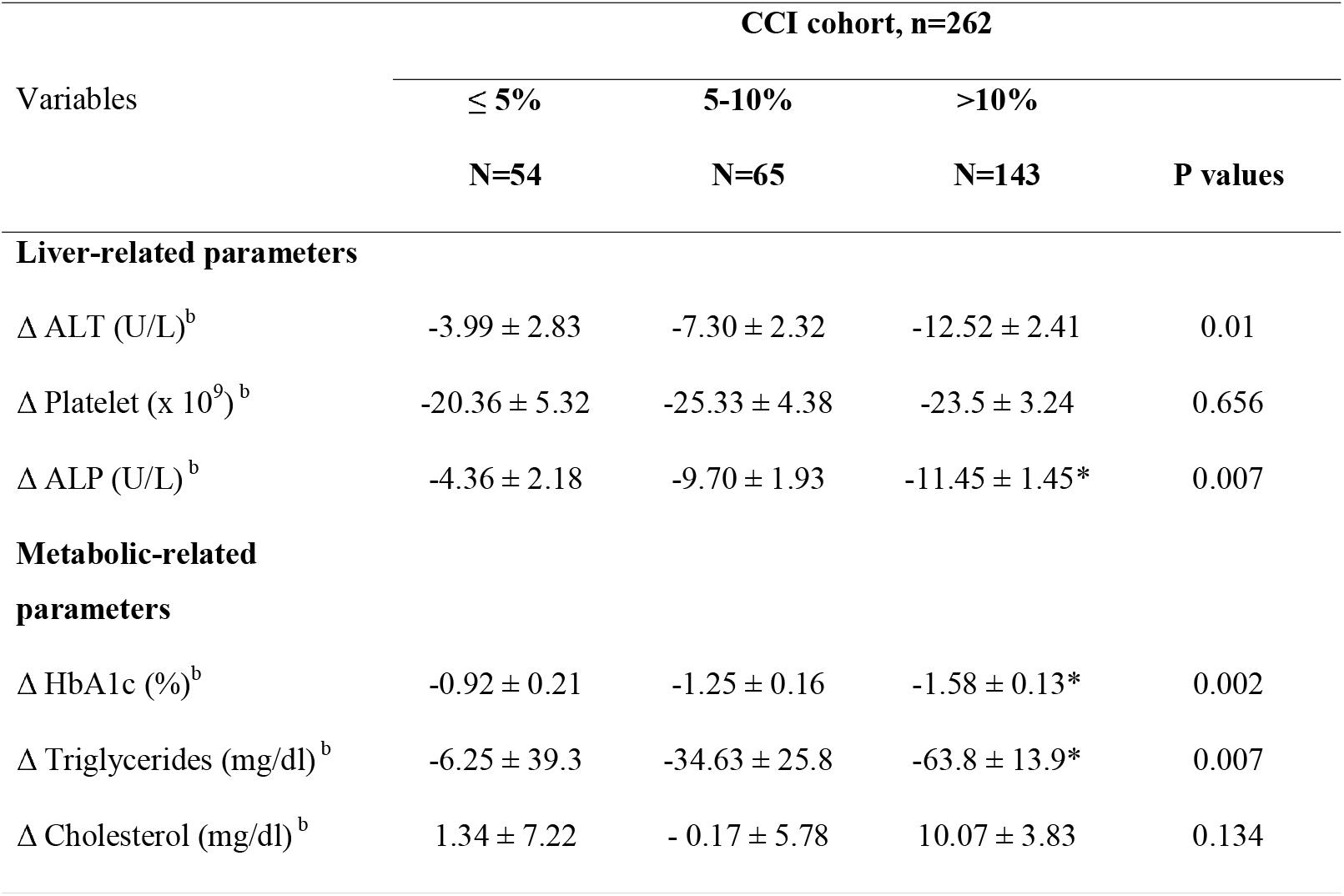
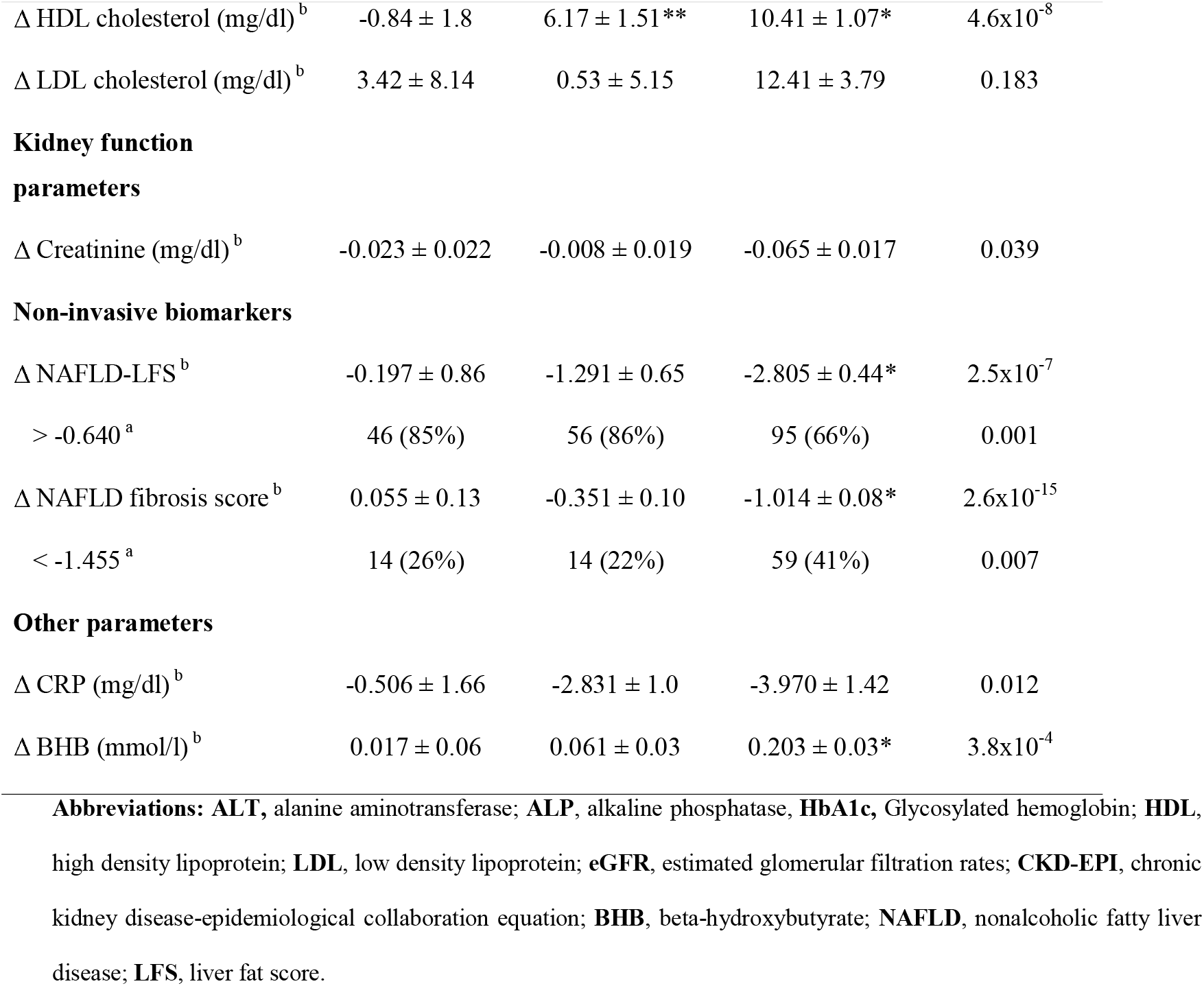
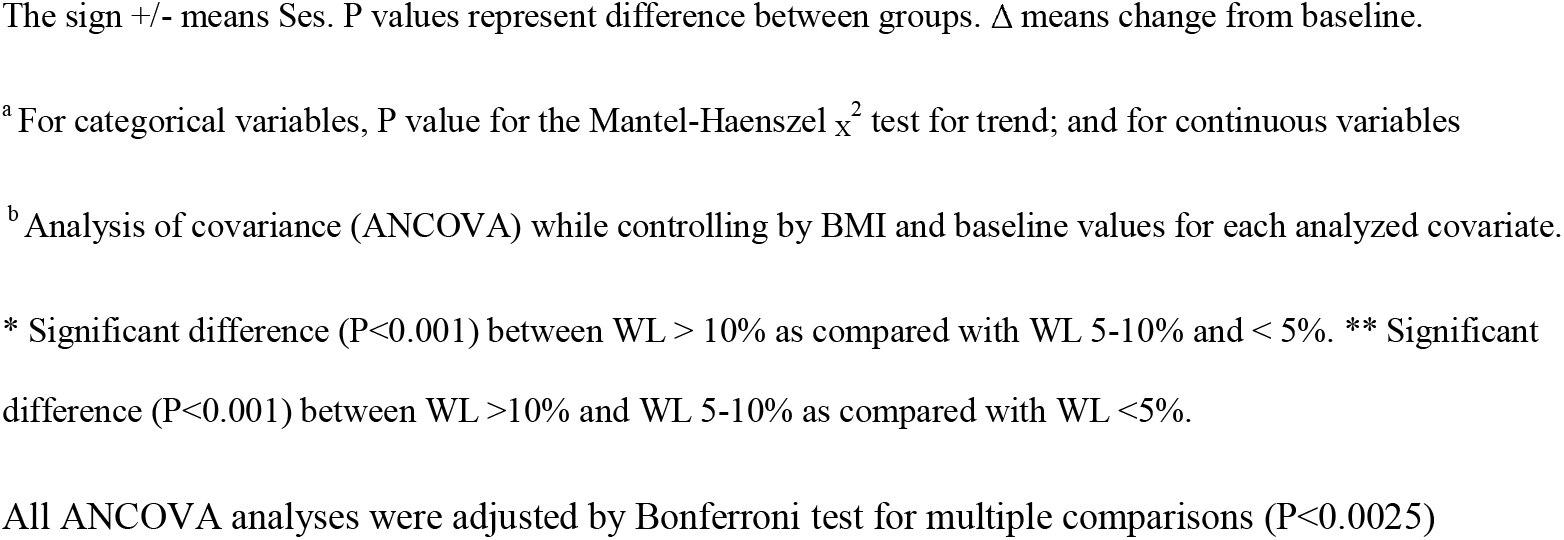
One-year associations between weight loss (%) and changes in liver- and diabetes-related variables. Intention-to-treat analysis.

**Table 4.**
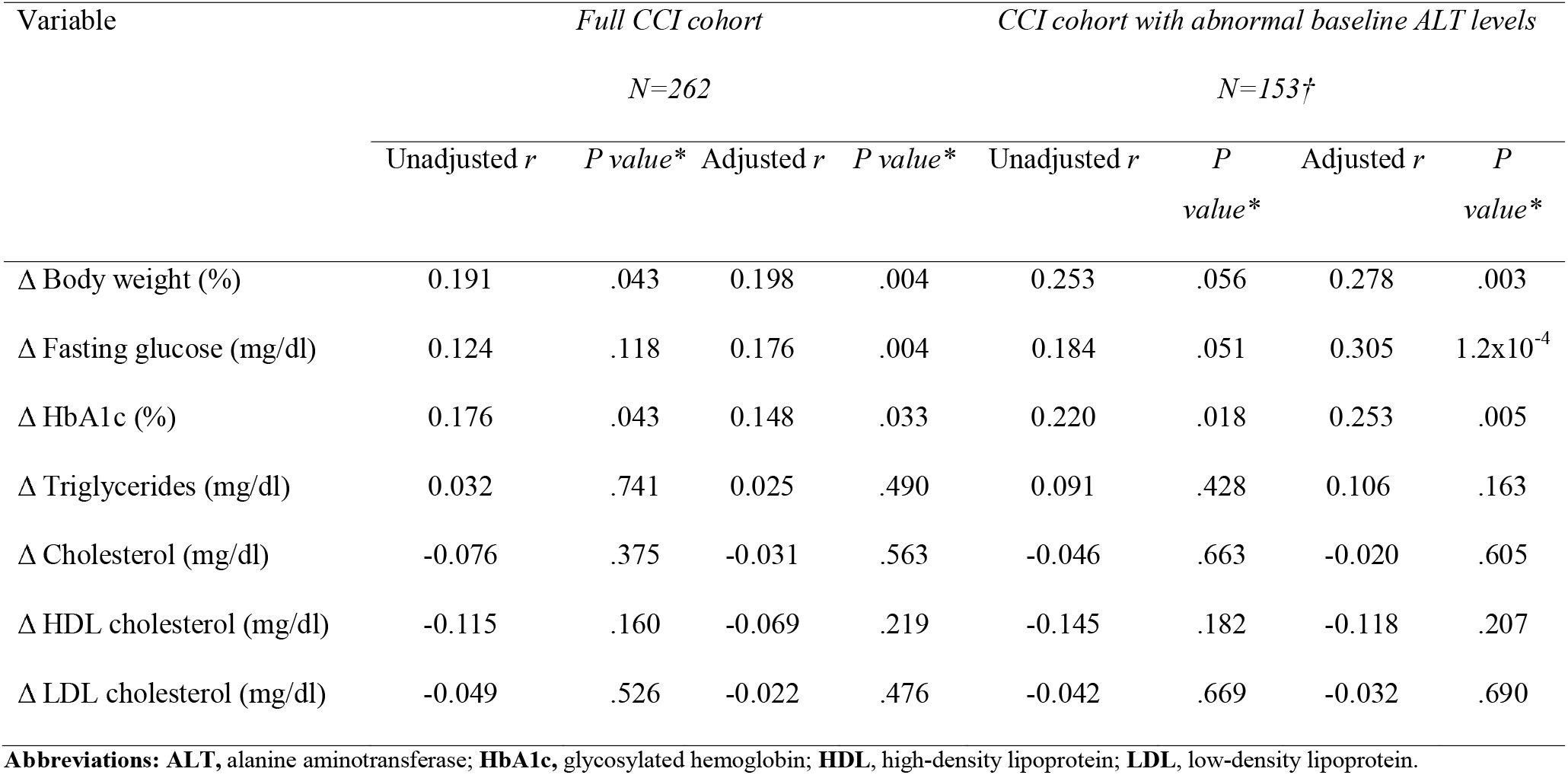
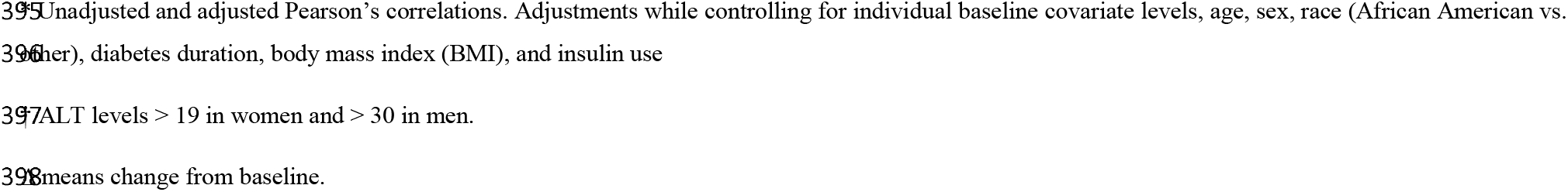
Correlations* change in ALT and changes in metabolic parameters.

## DISCUSSION

The findings of the current analysis show that one year of a digitally-supported CCI reduced risk of fatty liver and advanced liver fibrosis in overweight and obese adults with T2D. Improvements were concurrent with improved glycemic status, reduction in cardiovascular risk factors and decreased use of medications for diabetes and hypertension[36,47]. The beneficial effects extended to patients with increased levels of aminotransferase, thus indicating that remote care medically-supervised ketosis is also effective in patients at risk of liver disease progression. The influence of carbohydrate restriction and nutritional ketosis on liver histology of patients with biopsy-proven NASH remains largely unexplored in the context of a well-designed RCT. A pilot study including five patients with biopsy-proven NASH showed that 6-months of KD (less than 20 grams per day of carbohydrate) induced significant WL (mean of 13 kg) and four of five patients reduced liver fat, inflammation, and fibrosis [33]. The current study provides evidence that a remote-care medically-supervised KD can improve NASH and even fibrosis. A recent meta-analysis of ten studies reported the effects of LCD on liver function tests in patients with NAFLD, and concluded that LCD reduced IHLC, but did not improve liver enzymes[35], although heterogeneity among NAFLD populations and interventions were observed across the included studies.

Among CCI participants, correlations were also found between the improvements in HbA1c and ALT changes, even after controlling for WL and changes in insulin use. Among subjects with abnormal ALT levels at baseline, a reduction of ≥ 0.5% in HbA1c was associated with increased rates of ALT normalization. This finding suggests that liver enzyme improvements may be related to improvements in glycemic control and insulin concentration in addition to weight loss. Importantly, few studies have directly compared the metabolic advantages of different diets for the treatment of NAFLD[15, 32, 48], and the impact of dietary macronutrient composition remains largely unknown. Three studies have shown that low-carbohydrate and low-fat diets reduced liver fat, transaminases and insulin resistance to similar degrees[15, 21, 48], whereas another study reported that a moderate hypocaloric low-carbohydrate diet in insulin resistant patients improved ALT levels more than a hypocaloric low-fat diet, despite equal weight loss[48]. Among patients with T2D, a “moderate-carbohydrate modified Mediterranean diet” (35% carbohydrates, 45% high monounsaturated fat) showed greater ALT reductions than two other higher carbohydrate hypocaloric diets including the 2003 recommended ADA or low glycemic index diets[49].

Our results also demonstrated that non-invasive risk scores for fatty liver and fibrosis were improved in patients who underwent CCI as compared to the UC control, and greater reductions were observed in patients with the largest reductions in body weight (≥ 10%). Our results are consistent with previous studies reporting that LCD reduce intrahepatic lipid accumulation[15, 16, 21, 32, 33]. Likewise, 1-year liver fibrosis as assessed by NFS improved in the CCI group, and the proportion of patients with low likelihood of fibrosis increased from 18% to 33% at 1 year of intervention. Similar to previous studies addressing the impact of weight loss on NASH-related fibrosis[13, 50], we showed a relationship between the degree of weight loss and improvements in NFS.

LCD or KD have been proposed to more effectively reduce all features of the metabolic syndrome, which is present in approximately 80% of NAFLD patients, compared to low-fat diets [51, 52]; however, the physiological mechanisms are not fully established[53–55]. In line with our findings, Holland et al.[56] showed that irrespective of physical exercise, rats fed a ketogenic formulation had lower liver triglycerides and lower activation of the pro-inflammatory NF-kB pathway compared to rats fed Western and standard chow diets. Likewise, a recent human study using a two-week isocaloric carbohydrate restricted diet, not only demonstrated a drastic reduction of hepatic steatosis, but a shift in lipid metabolism pathway from de-novo lipogenesis to ß-oxidation and increased BHB production[57]. This shift in the lipid homeostasis following a short-term ketogenic diet occurred in conjunction with a shift in gut microbia towards increased folate production as well as decreased expression of key serum inflammatory markers[57].

Strengths and weaknesses of this clinical trial have been previously described[36]. Some strengths of this study include a large cohort of patients with T2D and high suspicion of NAFLD, an intervention with one-year of digitally-supported continuous care including monitored adherence to nutritional ketosis, and a control group of patients with T2D provided usual care with standard nutritional recommendations[36]. Relative to prior outpatient interventions, the current study is unusual in the degree of health coach and physician support, the degree of prescribed carbohydrate restriction and the use of BHB as a blood biomarker of dietary adherence. These attributes may contribute to superior outcomes observed in the intervention group when compared to UC patients. The multi-component approach used in the intervention, not only encouraged the patient to adapt carbohydrate restriction through continuous monitoring of nutritional ketosis but also provided behavioral support through interaction with their health coaches.

Some weaknesses of this study include the absence of imaging- or biopsy-proven NAFLD or NASH diagnosis and lack of random allocation to assign patients to intervention and control groups. Food was not provided for participants so dietary macronutrient and micronutrient contents and sources were not strictly controlled.

In conclusion, one year of a digitally-supported continuous care intervention including individualized nutritional ketosis led to significant improvement in non-invasive markers of liver fat and fibrosis together with sustained weight loss in overweight and obese type 2 diabetes patients. A relationship was observed between the degree of weight loss and improvements in liver- and non-liver-related outcomes with greater benefits in patients losing more than 10% of body weight. A reduction of ≥ 0.5% in HbA1c was independently associated with ALT normalization even after controlling for weight loss. Medical interventions incorporating ketogenic diets appear effective for improving NAFLD, and therefore, may be an effective approach for reversing the natural history of NAFLD progression, although further studies are needed to confirm potential beneficial effect in patients with biopsy-confirmed NASH.

## Supporting information

## Acknowledgments

Authors thank Drs. Marwan Ghabril and Raj Vuppalanchi for their helpful discussions with various analyses. The authors also thank the study participants for their active involvement in the study.

## Author contributions

E.V.G, S.J.A, R.N.A, J.P.M and N.P.C wrote the manuscript. A.L.M, N.H.B, S.J.H and S.J.A participated in data acquisition. E.V.G and S.J.A analyzed the data. N.P.C, S.J.H, N.H.B, A.L.M, W.W.C, J.P.M, S.D.P and J.S.V supervised this particular analysis and edited the manuscript. All authors approved the final version of the manuscript.

## Abbreviations

CCI: continuous care intervention;
UC: usual care;
NAFLD: non-alcoholic fatty liver disease;;
N-LFS: non-alcoholic liver fatty score;
NFS: non-alcoholic fatty liver disease fibrosis score;
CLD: chronic liver disease;
HCC: hepatocellular carcinoma;
T2D: type 2 diabetes;
NASH: nonalcoholic steatohepatitis;
BMI: body mass index;
LCHF: low-carbohydrate high-fat;
ALT: alanine aminotransferase;
AST: aspartate aminotransferase;
IHLC: intrahepatic lipid content;
ADA: American Diabetes Association;
BHB: beta-hydroxybutyrate;
CLIA: clinical laboratory improvement amendments;
ITT: intention-to-treat;
MIM: multiple imputation methods;
WL: weight loss;
EOT: end-of-treatment;
HDL: high density lipoprotein;
RCT: randomized controlled trial;
LCD: low-carbohydrate diet;
KD: ketogenic diet;
HFD: high-fat diet

## REFERENCES

1. Estes C, Razavi H, Loomba R, et al. Modeling the epidemic of nonalcoholic fatty liver disease demonstrates an exponential increase in burden of disease. Hepatology 2018;67:123–33.

2. Younossi ZM, Koenig AB, Abdelatif D, et al. Global epidemiology of nonalcoholic fatty liver disease-Meta-analytic assessment of prevalence, incidence, and outcomes. Hepatology 2016;64:73–84.

3. Byrne CD, Targher G. NAFLD: a multisystem disease. J Hepatol 2015;62:S47–64.

4. Adams LA, Anstee QM, Tilg H, et al. Non-alcoholic fatty liver disease and its relationship with cardiovascular disease and other extrahepatic diseases. Gut 2017;66:1138–53.

5. Portillo-Sanchez P, Bril F, Maximos M, et al. High Prevalence of Nonalcoholic Fatty Liver Disease in Patients With Type 2 Diabetes Mellitus and Normal Plasma Aminotransferase Levels. J Clin Endocrinol Metab 2015;100:2231–8.

6. Hazlehurst JM, Woods C, Marjot T, et al. Non-alcoholic fatty liver disease and diabetes. Metabolism 2016;65:1096–108.

7. Angulo P, Kleiner DE, Dam-Larsen S, et al. Liver Fibrosis, but No Other Histologic Features, Is Associated With Long-term Outcomes of Patients With Nonalcoholic Fatty Liver Disease. Gastroenterology 2015;149:389–97.

8. Ekstedt M, Hagstrom H, Nasr P, et al. Fibrosis Stage Is the Strongest Predictor for Disease-Specific Mortality in NAFLD After Up to 33 Years of Follow-Up. Hepatology 2015;61:1547–54.

9. Adams LA, Lymp JF, St Sauver J, et al. The natural history of nonalcoholic fatty liver disease: a population-based cohort study. Gastroenterology 2005;129:113–21.

10. Vilar-Gomez E, Calzadilla-Bertot L, Wai-Sun Wong V, et al. Fibrosis severity as a determinant of cause-specific mortality in patients with advanced nonalcoholic fatty liver disease: A multi-national cohort study. Gastroenterology 2018; 155: 443–57.

11. Chalasani N, Younossi Z, Lavine JE, et al. The diagnosis and management of nonalcoholic fatty liver disease: Practice guidance from the American Association for the Study of Liver Diseases. Hepatology 2018;67:328–57.

12. European Association for the Study of the L, European Association for the Study of D, European Association for the Study of O. EASL-EASD-EASO Clinical Practice Guidelines for the management of non-alcoholic fatty liver disease. J Hepatol 2016;64:1388–402.

13. Vilar-Gomez E, Martinez-Perez Y, Calzadilla-Bertot L, et al. Weight Loss Through Lifestyle Modification Significantly Reduces Features of Nonalcoholic Steatohepatitis. Gastroenterology 2015;149:367–78.

14. Zelber-Sagi S, Kessler A, Brazowsky E, et al. A double-blind randomized placebo-controlled trial of orlistat for the treatment of nonalcoholic fatty liver disease. Clin Gastroenterol Hepatol 2006;4:639–44.

15. Haufe S, Engeli S, Kast P, et al. Randomized Comparison of Reduced Fat and Reduced Carbohydrate Hypocaloric Diets on Intrahepatic Fat in Overweight and Obese Human Subjects. Hepatology 2011;53:1504–14.

16. Browning JD, Baker JA, Rogers T, et al. Short-term weight loss and hepatic triglyceride reduction: evidence of a metabolic advantage with dietary carbohydrate restriction. Am J Clin Nutr 2011;93:1048–52.

17. Ryan MC, Itsiopoulos C, Thodis T, et al. The Mediterranean diet improves hepatic steatosis and insulin sensitivity in individuals with non-alcoholic fatty liver disease. J Hepatol 2013;59:138–43.

18. Eckard C, Cole R, Lockwood J, et al. Prospective histopathologic evaluation of lifestyle modification in nonalcoholic fatty liver disease: a randomized trial. Therap Adv Gastroenterol 2013;6:249–59.

19. Wong VW, Chan RS, Wong GL, et al. Community-based lifestyle modification programme for non-alcoholic fatty liver disease: a randomized controlled trial. J Hepatol 2013;59:536–42.

20. Trovato FM, Catalano D, Martines GF, et al. Mediterranean diet and non-alcoholic fatty liver disease: the need of extended and comprehensive interventions. Clin Nutr 2015;34:86–8.

21. Kirk E, Reeds DN, Finck BN, et al. Dietary Fat and Carbohydrates Differentially Alter Insulin Sensitivity During Caloric Restriction. Gastroenterology 2009;136:1552–60.

22. Promrat K, Kleiner DE, Niemeier HM, et al. Randomized Controlled Trial Testing the Effects of Weight Loss on Nonalcoholic Steatohepatitis. Hepatology 2010;51:121–9.

23. Tobias DK, Chen M, Manson JE, et al. Effect of low-fat diet interventions versus other diet interventions on long-term weight change in adults: a systematic review and meta-analysis. Lancet Diabetes Endocrinol 2015;3:968–79.

24. Sackner-Bernstein J, Kanter D, Kaul S. Dietary Intervention for Overweight and Obese Adults: Comparison of Low-Carbohydrate and Low-Fat Diets. A Meta-Analysis. Plos One 2015;10.

25. Mansoor N, Vinknes KJ, Veierod MB, et al. Effects of low-carbohydrate diets v. low-fat diets on body weight and cardiovascular risk factors: a meta-analysis of randomised controlled trials. Br J Nutr 2016;115:466–79.

26. Gardner CD, Kiazand A, Alhassan S, et al. Comparison of the Atkins, Zone, Ornish, and LEARN diets for change in weight and related risk factors among overweight premenopausal women: the A TO Z Weight Loss Study: a randomized trial. JAMA 2007;297:969–77.

27. Goday A, Bellido D, Sajoux I, et al. Short-term safety, tolerability and efficacy of a very low-calorie-ketogenic diet interventional weight loss program versus hypocaloric diet in patients with type 2 diabetes mellitus. Nutr Diabetes 2016;6: e230. doi: 10.1038/nutd.2016.36.

28. Wheeler ML, Dunbar SA, Jaacks LM, et al. Macronutrients, Food Groups, and Eating Patterns in the Management of Diabetes A systematic review of the literature, 2010. Diabetes Care 2012;35:434–445.

29. Johnstone AM, Horgan GW, Murison SD, et al. Effects of a high-protein ketogenic diet on hunger, appetite, and weight loss in obese men feeding ad libitum. Am J Clin Nutr 2008;87:44–55.

30. Volynets V, Machann J, Kuper MA, et al. A moderate weight reduction through dietary intervention decreases hepatic fat content in patients with non-alcoholic fatty liver disease (NAFLD): a pilot study. Eur J Nutr 2013;52:527–35.

31. Kani AH, Alavian SM, Esmaillzadeh A, et al. Effects of a novel therapeutic diet on liver enzymes and coagulating factors in patients with non-alcoholic fatty liver disease: A parallel randomized trial. Nutrition 2014;30:814–21.

32. de Luis DA, Aller R, Izaola O, et al. Effect of two different hypocaloric diets in transaminases and insulin resistance in nonalcoholic fatty liver disease and obese patients. Nutr Hosp 2010;25:730–5.

33. Tendler D, Lin SY, Yancy WS, et al. The effect of a low-carbohydrate, ketogenic diet on nonalcoholic fatty liver disease: A pilot study. Dig Dis Sci 2007;52:589–93.

34. Huang MA, Greenson JK, Chao C, et al. One-year intense nutritional counseling results in histological improvement in patients with non-alcoholic steatohepatitis: A pilot study. Am J Gastroenterol 2005;100:1072–81.

35. Haghighatdoost F, Salehi-Abargouei A, Surkan PJ, et al. The effects of low carbohydrate diets on liver function tests in nonalcoholic fatty liver disease: A systematic review and meta-analysis of clinical trials. J Res Med Sci 2016;21:53.

36. Hallberg SJ, McKenzie AL, Williams PT, et al. Effectiveness and safety of a novel care model for the management of type 2 diabetes at 1 year: an open-label, non-randomized, controlled study. Diabetes Ther 2018; 9:583–612.

37. Standards of Medical Care in Diabetes-2015: Summary of Revisions. Diabetes Care 2015;38:S4–S4.

38. Kotronen A, Peltonen M, Hakkarainen A, et al. Prediction of Non-Alcoholic Fatty Liver Disease and Liver Fat Using Metabolic and Genetic Factors. Gastroenterology 2009;137:865–872.

39. Machado MV, Cortez-Pinto H. Non-invasive diagnosis of non-alcoholic fatty liver disease. A critical appraisal. J Hepatol 2013;58:1007–19.

40. Angulo P, Hui JM, Marchesini G, et al. The NAFLD fibrosis score: a noninvasive system that identifies liver fibrosis in patients with NAFLD. Hepatology 2007;45:846–54.

41. Vilar-Gomez E, Chalasani N. Non-invasive assessment of non-alcoholic fatty liver disease: Clinical prediction rules and blood-based biomarkers. J Hepatol 2018;68:305–15.

42. Sun W, Cui H, Li N, et al. Comparison of FIB-4 index, NAFLD fibrosis score and BARD score for prediction of advanced fibrosis in adult patients with non-alcoholic fatty liver disease: A meta-analysis study. Hepatol Res 2016; 46:862–70.

43. Prati D, Taioli E, Zanella A, et al. Updated Definitions of Healthy Ranges for Serum Alanine Aminotransferase Levels. Ann Intern Med. 2002;137:1–10.

44. Kline RB. Convergence if structural equation modeling and multilevel modeling. In M. Williams & W.P. Vogt (Eds.), Handbook of methodological innovation in social research methods 2011; 562–589.

45. John JA, Draper NR. An alternative family of transformations. Appl Statist 1980; 29: 190–197.

46. Kim WR, Flamm SL, Di Bisceglie AD, et al. Serum activity of alanine aminotransferase (ALT) as an indicator of health and disease. Hepatology 2008; 47: 1363–70.

47. Bhanpuri NH, Hallberg SJ, Williams PT, et al. Cardiovascular Disease Risk Factor Responses to a Type 2 Diabetes Care Model Including Nutritional Ketosis Induced by Sustained Carbohydrate Restriction at One Year: An Open Label, Non-Randomized, Controlled Study (in press).

48. Ryan MC, Carter S, Abbasi F, et al. Serum alanine aminotransferase levels decrease further with carbohydrate than fat restriction in insulin-resistant adults. Diabetes Care 2007;30:1075–80.

49. Fraser A, Abel R, Lawlor DA, et al. A modified Mediterranean diet is associated with the greatest reduction in alanine aminotransferase levels in obese type 2 diabetes patients: results of a quasi-randomised controlled trial. Diabetologia 2008;51:1616–22.

50. Glass LM, Dickson RC, Anderson JC, et al. Total body weight loss of >/= 10 % is associated with improved hepatic fibrosis in patients with nonalcoholic steatohepatitis. Dig Dis Sci 2015;60:1024–30.

51. Volek JS, Fernandez ML, Feinman RD, et al. Dietary carbohydrate restriction induces a unique metabolic state positively affecting atherogenic dyslipidemia, fatty acid partitioning, and metabolic syndrome. Prog Lipid Res 2008;47:307–18.

52. Volek JS, Phinney SD, Forsythe CE, et al. Carbohydrate Restriction has a More Favorable Impact on the Metabolic Syndrome than a Low Fat Diet. Lipids 2009;44:297–309.

53. Samaha FF, Iqbal N, Seshadri P, et al. A low-carbohydrate as compared with a low-fat diet in severe obesity. N Engl J Med 2003;348:2074–81.

54. Foster GD, Wyatt HR, Hill JO, et al. A randomized trial of a low-carbohydrate diet for obesity. N Engl J Med 2003;348:2082–90.

55. Yancy WS, Jr., Olsen MK, Guyton JR, et al. A low-carbohydrate, ketogenic diet versus a low-fat diet to treat obesity and hyperlipidemia: a randomized, controlled trial. Ann Intern Med 2004;140:769–77.

56. Holland AM, Kephart WC, Mumford PW, et al. Effects of a ketogenic diet on adipose tissue, liver, and serum biomarkers in sedentary rats and rats that exercised via resisted voluntary wheel running. Am J Physiol Regul Integr Comp Physiol 2016;311:R337–51.

57. Mardinoglu A, Wu H, Bjornson E, et al. An integrated understanding of the rapid metabolic benefits of a carbohydrate-restricted diet on hepatic steatosis in humans. Cell Metab 2018; https://doi.org/10.1016/j.cmet.2018.01.005.

