## Supplementary material for "Post-hoc analyses of Surrogate Markers of Non-Alcoholic Fatty Liver Disease (NAFLD) and Liver Fibrosis in Patients with Type 2 Diabetes in a Digitally-Supported Continuous Care Intervention: An Open Label, Non-Randomized, Controlled Study"

**SUPPLEMENTARY APPENDIX**

**TABLE OF CONTENTS**

1. Methods……………...Page 2
2. Tables…………..........Page 4
3. Figures Legend……....Page 9

**METHODS**

***Study design and participants***

Briefly, this was a non-randomized and open-label controlled longitudinal study, including patients between 21 to 65 years old with a diagnosis of T2D and a BMI > 25 kg/m^2^. Patients were initially screened for the study based on the study’s inclusion and exclusion criteria[35]. Major exclusion criteria included serious renal, or cardiovascular dysfunction, hepatic failure, infectious disease, uncontrolled psychiatric disorder, history of ketoacidosis, a intolerance to dietary fat, cancer with active treatment in the last five years, and pregnancy or planned pregnancy. Further, patients with high alcohol intake defined as average consumption of 3 or more alcohol-containing beverages daily or consumption of more than 14 standard drinks per week were excluded. Patients on CCI had access to a remote care team consisting of a personal health coach and medical providers (physician or nurse practitioner). The participants in the CCI self-selected between two different educational modes; either via on-site education classes (n=136, CCI-onsite) or via web-based educational contents (n=126, CCI-virtual). The CCI patients were routinely assessed for nutritional ketosis based on blood beta-hydroxybutyrate (BHB) concentrations. The on-site and virtual patients were grouped together for analyses since no significant differences were observed in biochemical markers between these two modes of educational delivery[35]. We also recruited and followed a cohort of patients with T2D (n=87) who were categorized as UC[35]. This group of patients received a standard diabetes care treatment from their primary care physician or endocrinologist without modification. These patients were aware of the intervention cohort and could participate in that group if they chose.

***Interventions***

*CCI including personalized nutrition*

The CCI included support from a medical provider and health coach, education in nutrition and behavior change, peer support and individualized advice for maintaining nutritional ketosis during 1 year as described[35]. Briefly, all subjects were instructed to follow a ketogenic diet incorporating their personal preferences; health coaches monitored glycemic and ketosis status through patient reported daily blood glucose and blood BHB tests with a BHB target range of 0.5-3.0 mmol/L. Patients’ dietary modifications included restricting total dietary carbohydrate to a target of less than 30 g daily. Daily protein intake was targeted to 1.5 g/kg of reference body weight. Patients were encouraged to consume dietary fat to satiety, by consuming adequate omega-3 and omega-6 polyunsaturated fatty acids with the remaining fats consumed coming from monounsaturated and saturated fatty acids. Patients were also counseled on adequate intake of minerals, fluids and non-starchy vegetables[35].

*Usual care (UC)*

Usual care for these participants was continued by their own primary care physician (PCP) or endocrinologist, and registered dietitians counseled UC participants on diabetes self-management, nutrition, and lifestyle based on the American Diabetes Association (ADA) recommendations[37].

| **Score** | **Equation** |
| --- | --- |
| NAFLD liver fat score (N-LFS) | -2.89 + 1.18 x **metabolic syndrome** (yes=1 or no=0) + 0.45 x **type 2 diabetes** (yes=2 or no=0)* + 0.15 x **fasting insulin** (mU/l) + 0.04 x **fasting serum AST** (U/L) – 0.94 x **AST/ALT** |
| NAFLD fibrosis score (NFS) | −1.675 + 0.037 × **Age** (yrs) + 0.094 × **BMI** (kg/m^2^) + 1.13 × **IFG/diabetes** (yes = 1, no = 0) + 0.99 × **AST/ALT ratio** − 0.013 × **Platelet** (×10^9^/L) −0.66 × **Albumin** (g/dl) |

**Equations for calculating NAFLD liver fat score and NAFLD fibrosis score.**

**TABLES**

**Supplemental Table 1. Impact of CCI on weight loss based on BMI classes at baseline.**

|  | **BMI (kg/m^2^) at baseline** | | | | |
| --- | --- | --- | --- | --- | --- |
|  | **25-29.9**  **N=22** | **30-34.9**  **N=50** | **35-35.9**  **N=70** | **>40**  **N=120** | **P value** |
|  |  |  |  |  | 0.121† |
| <5%, n (%) | 6 (33) | 14 (29) | 16 (17) | 18 (18) |  |
| 5-10%, n (%) | 5 (24) | 14 (27) | 17 (29) | 30 (33) |  |
| ≥10%, n (%) | 11 (43) | 22 (43) | 37 (54) | 72 (59) |  |

† Mantel-Haenszel chi-square test for overall trend.

The +/- sign means SEs.

**FIGURE LEGENDS**

**Supplemental Figure 1.** Flow of patients through the study. Final analyses were performed on imputed data generated using a model of multiple imputation. Patients “Assessed for eligibility” are those patients who successfully screened for eligibility through phone conversation.

**Supplemental Figure 2**. Weight loss (%) at 1 year of intervention.

Weight loss (%) categories and stratification of patients in each category by treatment, UC and CCI based on ITT analysis

**Supplemental Figure 1**

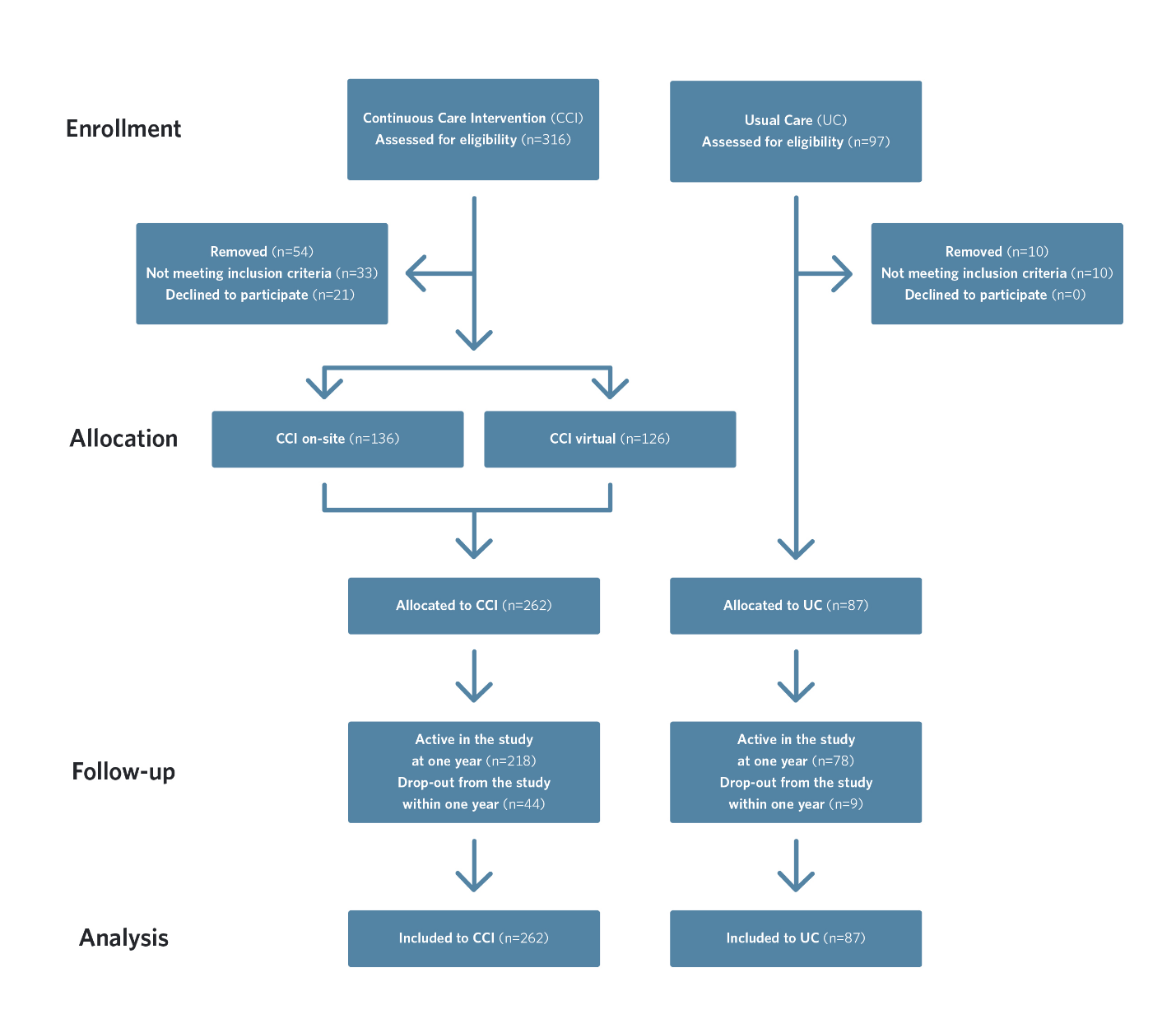

**Supplemental Figure 2**

P<0.001

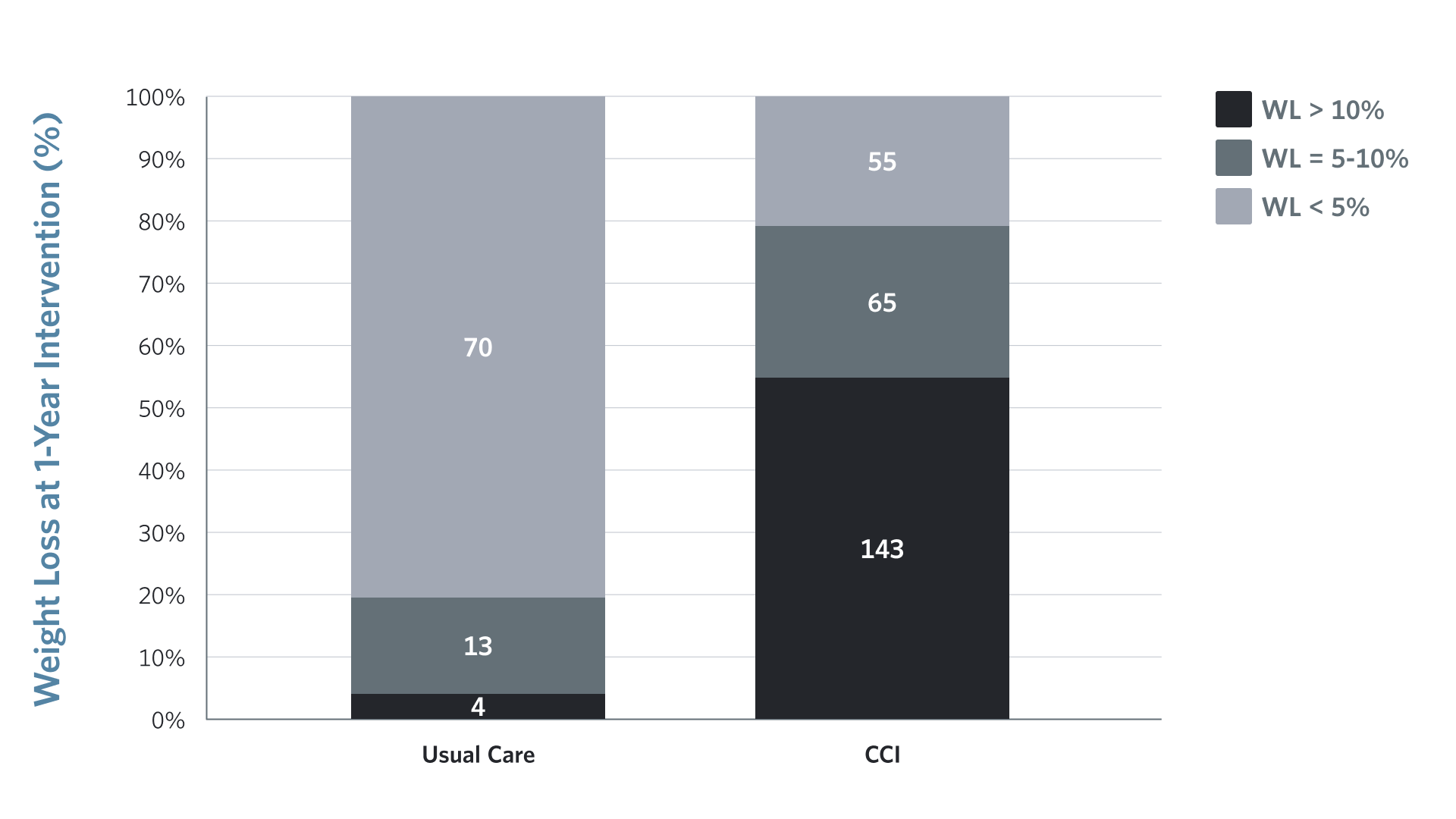
